# Bcl6 Preserves the Suppressive Function of Regulatory T Cells during Tumorigenesis

**DOI:** 10.1101/2020.01.21.914812

**Authors:** Yiding Li, Zhiming Wang, Huayu Lin, Lisha Wang, Xiangyu Chen, Qiao Liu, Qianfei Zuo, Jianjun Hu, Haoqiang Wang, Junyi Guo, Luoyingzi Xie, Jianfang Tang, Zhirong Li, Li Hu, Lilin Ye, Qizhao Huang, Lifan Xu

**Author notes:** **Correspondence:** Lifan Xu, Qizhao Huang, Lilin Ye.

## Abstract

During tumorigenesis, tumor infiltrating regulatory T (Treg) cells restrict the function of effector T cells in tumor microenvironment and thereby promoting tumor growth. The anti-tumor activity of effector T cells can be therapeutically unleashed, and is now being exploited for the treatment of various types of human cancers. However, the immune suppressive function of Treg cells remains a major hurdle to broader effectiveness of tumor immunotherapy. In this article, we reported that the deletion of Bcl6 specifically in Treg cells led to stunted tumor growth, which was caused by impaired Treg cell responses. Notably, Bcl6 is essential in maintaining the lineage stability of Treg cells in tumor microenvironment. Meanwhile, we found that the absence of follicular regulatory T (Tfr) cells, which is a result of Bcl6 deletion in Foxp3^+^ cells, was dispensable for tumor control. Importantly, the increased Bcl6 expression in Treg cells is associated with poor prognosis of human colorectal cancer and lymph node metastasis of skin melanoma. Furthermore, Bcl6 deletion in Treg cells exhibits synergistic effects with immune checkpoint blockade therapy. Collectively, these results indicate that Bcl6 actively participates in regulating Treg cell immune responses during tumorigenesis and can be exploited as a therapeutic target of anti-tumor immunity.

## 1 Introduction

Regulatory T (Treg) cells, with the master regulator Foxp3 (forkhead box P3), represent a functional distinct subset of CD4 T cells, which are endowed with the ability of immune suppression and play a pivotal role in maintaining immune-homeostasis and autoimmune diseases (1–3). Foxp3^+^ Treg cells exert their effector functions through a variety of molecular mechanisms. Firstly, Treg cells constitutively express the high-affinity heterotrimeric interleukin 2 (IL2) receptor, also known as CD25, which further bind to and consume IL2 from their surroundings, thus compromising its effects on non-Foxp3 effector T cells (Teff) (4,5). Treg cells also express high level of cytotoxic T lymphocyte antigen 4 (CTLA4), which can bind to CD80/CD86 on antigen presenting cells (APCs) and thereby transmitting suppressive signals to these cells and reducing their capacity to activate Teff cells (6). Besides, CTLA4 exhibits a higher affinity to CD80/CD86 than that of CD28, thus competing with this co-stimulatory receptor, which further disrupts the priming and/or activation of Teff cells (7). Additionally, Treg cells can produce immunosuppressive cytokines, such as TGFβ, IL10, and IL35 (8), which will downregulate the activity of APCs and Teff cells, and secrete granzymes and perforin that can directly kill these cells (9).

Akin to CD8 memory T cells consisting of effector memory T cells (TEM), central memory T cells (TCM) and tissue-resident memory T cells (TRM) (10), thymus derived Treg (tTreg) cells can also be divided as central (cTreg) and effector (eTreg) Treg cells based on the expression of trafficking receptors (11). cTreg cells are programmed to recirculate through secondary lymphoid organs (SLOs) by expressing CD62L as well as CCR7, while eTreg cells capable of entering non-lymphoid tissues by virtue of expressing chemokine receptors such as CXCR3, CCR4, CCR6, CCR2, and CCR5, etc. (12) Adoptive transfer studies have shown that cTreg cells could convert into more highly proliferative eTreg cells in response to tissue self-antigens (13). Notably, eTreg cells have also been observed in increased numbers within diverse experimental mouse tumors and human cancers which suggest the involvement of Treg cells in anti-tumor immunity (14,15).

The depletion of Treg cells, which results in tumor rejection and retardation of tumor growth, has been reported in mouse models (16–18). For several types of human cancers, including melanoma, non-small-cell lung, gastric and ovarian cancers, Treg cells account for 10–50% of tumor infiltrated CD4 T cells compared with 2–5% of CD4 T cells in the peripheral blood of healthy individuals (19,20). Furthermore, a relatively high infiltration abundance of Treg cells versus non-Treg cells in TME is associated with poor prognosis in cancer patients (21). Using single cell sequencing, it has been found that the majority of tumor infiltrating Treg cells are uniformly highly activated comparing with those in periphery tissues (22), with lower level of CD45RA and CCR7 expression, and highly enriched for a range of co-stimulatory molecules such as CD27, ICOS, OX40, 41BB, GITR, and co-inhibitory molecules such as CTLA4, PD1, LAG3, and TIGIT. Remarkably, a number of cytokines and chemokine receptor genes, most notably CCR8, were upregulated in tumor-resident Treg cells in comparison to normal tissue-resident ones (23). These information supports the notion that tumor infiltrating Treg cells hold a distinct transcriptional profiling however, the underlying mechanisms regulating Treg cells within TME still remain obscure (22).

The transcription factor B cell lymphoma 6 (Bcl6) is intensively investigated during the development of several types of T cell subpopulations and its fine-tuned expression is modulated by a net of cytokines (e.g., IL6, IL12, IFNγ, type I IFN signaling, TGFβ, and TNFα) in a variety of cell types and repressed by IL2-STAT5 signaling (24–27). It controls T follicular helper (Tfh) cell development and is expressed by Foxp3^+^ T follicular regulatory (Tfr) cells (28–30). The specific deletion of Bcl6 in Treg cells in mouse models do not result in spontaneous inflammatory disease, however, it did enhance Th2-mediated airway inflammation following immunization (31). Bcl6-deficient Treg cells express higher level of GATA3 when compared with WT (wild-type) Treg cells, and it seems that Bcl6 controls the Th2 inflammatory activity of Treg cells by repressing GATA3 (32). We revealed that together with TCF1, Bcl6 is critical in transducing mTORC1 signaling in regulating Tfr differentiation during protein immunization or viral infection (33). However, whether Bcl6 possesses a role in regulating Treg cells during tumorigenesis has not been explored.

In this study, we found that Bcl6 expression is enhanced in Treg cells under TME, indicating that Bcl6 potentially serves to regulating the suppressive function of Treg cells on Teff cells in TME. Accordingly, depleting Bcl6 in Treg cells enhanced anti-tumor capability and retarded tumor progression. Moreover, this anti-tumor effect can synergize with anti-CTLA4 or anti-PD1 therapy, thus may serve as a therapeutic target to further improve the efficacy of immune checkpoint blockade responders.

## 2 Materials and methods

### 2.1 Mice

Bcl6-RFP (*Bcl6*^rfp/+^ or *Bcl6*^rfp/rfp^) mice were generously provided by Dr. Xindong Liu (Southwest Hospital, Chongqing, China). *Foxp3*^DTR^ were generously provided by Dr. Hua Tang (Institute of Immunology, Shandong First Medical University, Jinan, China). CXCR5-GFP knock-in mice have been described previously (34). *Bcl6*^fl/fl^, *Foxp3*^YFP-Cre^ knock-in, Cxcr5^−/−^, C57BL/6J (CD45.1 and CD45.2) mice were purchased from Jackson Laboratory. *Bcl6*^fl/fl^ mice were bred with *Foxp3*^YFP-Cre^ knock-in mice to generate *Bcl6*^fl/fl^*Foxp3*^Cre^ mice. All these strains are C57BL/6 background. All the mice used were analyzed at 6-10 weeks of age (indicated in diagram as “Sac”), and both genders were included without randomization or “binding”. Bone marrow (BM) chimeras were used after 8-10 weeks of reconstitution. LCMV virus (Armstrong strain) was provided by R. Ahmed (Emory University) and propagated in our laboratory as previously described (35). And 2×10^5^ plaque-forming units of this strain were used to establish acute infection in mice. For all the phenotypic analysis, at least 3 animals of each genotype with matched age and gender were analyzed.

### 2.2 Tissue Preparation

Spleens were surgically removed with sterilized surgical equipment and crushed with the blunt of 1mL syringe on Petri dishes containing 3mL of red blood cell lysis buffer. The spleen mixtures were separately filtered through a 70μM filter into a 15mL conical centrifuge tube, centrifuged at 1800rpm for 6min at 4°C. After wash, cell pellets were resuspended in 5mL of R2 media (RPMI-1640 (SIGMA Cat. RNBH7001) + 2% fetal bovine serum (FBS; gibco Cat. 10270-106)). Draining lymph nodes (dLNs) were extracted with sterilized surgical equipment and crushed between the frosted surfaces of super-frosted microscope slides into wells containing R2. Cell mixtures were then filtered through a 70μM filter into a 15mL conical centrifuge tube, centrifuged at 1800rpm for 6min at 4°C. After wash, cell pellets were resuspended in 0.5mL of R2 media. Tumors were removed from mice with sterile surgical instruments, pictured and weighted then shredded with ophthalmic scissors. Tumor tissue mixtures were transferred into 15mL conical tubes and filled with collagenase digest media (R2+Collagenase). B16-F10 Lung tumor tissue were treated with type2 collagenase (Sangon Biotech Cat. A004174-0001) and MC-38 solid tumor tissues were treated with type1 collagenase (Sangon Biotech Cat. A004194-0001). Samples were subsequently placed on a 37°C shaker for 1 hour, then filtered through 100μM filters into 50mL conical tubes and washed with R2 before centrifugation. B16-F10 tumor cells were further fractionated 2000rpm for 30 min at 4°C on a two-step gradient consisting 44% and 67% Percoll solutions (GE Cat. 17-0891-09). The T cell fraction was recovered from the inter-face between the 2 layers.

### 2.3 Flow Cytometry and Antibodies

Flow cytometry data were acquired with a FACSCantoⅡ (BD Biosciences) and were analyzed with FlowJo software (Tree Star). The antibodies and reagents used for flow cytometry staining are listed in Supplementary Table 1. Surface staining was performed in PBS containing 2% BSA or FBS (w/v). Tfh cell staining has been described (36). Staining of Bcl6, Bcl2, Tbet and Foxp3 were performed with the Foxp3/Transcription Factor Staining Buffer Set (00-5523; eBioscience). For in vivo incorporation of the thymidine analog BrdU, mice were given BrdU (1.5mg BrdU (5-bromodeoxyuridine) in 0.5ml PBS) intraperitoneally 3h before mice were sacrificed. BrdU in T cells was stained with a BrdU Flow Kit (552598; BD Biosciences) according to the manufacturer’s instructions. For the detection of cytokine production, lymphocytes were stimulated for 5h in the presence of PMA (50ng/ml), ionomycin (1μg/ml), Golgi Plug, Golgi Stop, anti-CD107a and anti-CD107b antibodies (BD Bioscience). Intracellular cytokine staining for CD107, granzyme B and Ki67 were performed with the Cytofix/Cytoperm Fixation/Permeabilization Kit (554714, BD Biosciences).

### 2.4 Adoptive Transfer and Generation of Bone Marrow Chimeras

A total of approximately 1×10^6^ splenocytes with WT Treg cells 1:1 mixed with KO Treg cells were adoptively transferred into cyclophosphamide (CTX, Sigma) treated (single dose at 200mg/kg, 12-24 hours before T cell transfer) CD4^−/−^ mice, which were inoculated with B16-F10 cells intravenously or MC-38 intraperitoneally on the following day. Bone marrow was collected from *Foxp3*^DTR^ and *Cxcr5*^−/−^ mice. Approximately 5×10^6^ cells of a 1:1 mixture of *Foxp3*^DTR^ and *Cxcr5*^−/−^ bone marrow cells were transferred intravenously into sub-lethally irradiated (two doses of 4.5 Gy each) CD4^−/−^ recipients. Recipient mice were reconstituted for 8-10 weeks before tumor transplantation.

### 2.5 Cell culture and In vitro suppression assay

Tumor cell lines, B16-F10, B-Luciferase B16-F10 (BIOCYTOGEN Cat. B-MCL-002) and MC-38 were cultured for cell injection into C57BL/6J mice. B16-F10, B-Luciferase B16-F10 and MC-38 cell medium was composed of DMEM (gibco Cat. C11995500BT), 10% FBS), 1% L-Glutamine (Solarbio Cat. G0200), and 1% penicillin/streptomycin (gibco Cat. 15070-063). Prior to injection, cells were adjusted with RPMI, each needle containing 100μL/1×10^6^ (in situ model) or 500μL/5×10^5^ tumor cells. (metastasis model).

CD4^+^CD25^+^GITR^+^CXCR5^−^ and CD4^+^CD25^+^GITR^+^CXCR5^+^ T cells from WT or *Bcl6*^fl/fl^*Foxp3*^Cre^ mice were sorted from tumor tissues or dLNs. CD8^+^CD44^+^ T cells from WT mice were sorted from dLNs. 100,000 CD8^+^ Teffs were incubated with different numbers of CD4^+^CD25^+^GITR^+^CXCR5^−^ T cells from tumor bearing WT, *Bcl6*^fl/fl^*Foxp3*^Cre^ or naive WT, *Bcl6*^fl/fl^*Foxp3*^Cre^ mice at 37°C for 72 hours and were assessed by flow cytometry.

### 2.6 Quantitative RT-PCR

For comparison of gene expression in *Bcl6*^−/−^ Treg cells and wild-type Treg cells both in naive state and tumor bearing state, Treg cells (CD4^+^CD25^+^GITR^+^CXCR5^−^) were sorted from naive or tumor implanted mice and were subsequently lysed in TRIzol LS reagent (10296; Life Technologies). Total RNA was extracted and reverse-transcribed with a RevertAid H Minus First Strand cDNA Synthesis Kit (K1632; Thermo Scientific). The resulting cDNA was analyzed for the expression of various target genes with a QuantiTest SYBR Green PCR Kit (204143; Qiagen) and the corresponding primers on a CFX96 Touch Real-Time System (Bio-Rad) (Supplementary Table 2).

### 2.7 Analysis of TCGA data

The TCGA normalized RNA-seq data of human colorectal cancer and skin melanoma patients were downloaded from https://genome-cancer.ucsc.edu/. None of the patients had any record of immunotherapy treatment. 352 human colorectal patients with primary tumor were included into analysis. Samples/patients were split based on gene expression levels analyzed by receiver operating characteristic curve and grouped into Foxp3^lo^ (66.76%) and Foxp3^hi^ (33.24%). Foxp3^hi^ ones were further divided into Bcl6^lo^ (50.64%) and Bcl6^hi^ (49.36%) subpopulations for analysis. Survival curves were then generated, and differences were evaluated by Log-rank test. 84 primary skin melanoma patients and 213 patients with lymph node metastasis were included. Samples/patients were split based on gene expression levels analyzed by receiver operating characteristic curve.

### 2.8 Statistical Analysis

Statistical analysis was conducted with Prism 7 software (GraphPad). An unpaired two-tailed t-test with 95% confidence interval was used to calculate P-values. For bone marrow and splenic chimera experiments, a paired two-tailed t-test with 95% confidence interval was used to calculate P values. Tumor growth curves at different time points were plotted by two-way ANOVA with a Turkey post hoc test comparison among groups.

## 3 Results

### 3.1 The deficiency of Bcl6 in Treg cells leads to delayed tumor progression

To investigate the role of Bcl6 in Treg cells immune response during tumorigenesis, we first evaluated the expression pattern of Bcl6 in Treg cells in TME and tumor dLNs of wild-type (WT) mice intravenously implanted with B16-F10 melanoma cells (metastasis model). Using real-time PCR, we observed an upregulation of Bcl6 transcripts in tumor infiltrating Treg cells when compared with dLNs derived Treg cells (TdLN Treg) and CD44^−^CD4^+^ T cells (naive CD4) (Figure 1A). Using flow cytometric analysis, we confirmed that tumor infiltrating Treg cells expressed higher level of Bcl6 than Treg cells derived from tumor dLNs and spleens (Figure 1B) in mice challenged with MC-38 cells subcutaneously (in situ model). The enhanced expression of Bcl6 in tumor infiltrating Treg cells indicates an important but unsolved role of Bcl6 within TME. To this end, we crossed *Bcl6*^fl/fl^ mice with *Foxp3*^YFP-Cre^ (*Foxp3*^Cre^) mice to generate mice in which the *Bcl6* alleles are conditionally deleted in Foxp3^+^ Treg cells (hereafter referred to as *Bcl6*^fl/fl^*Foxp3*^Cre^ mice). Then we challenged these mice with MC-38 cells or B16-F10 melanoma cells. We observed that tumor growth was significantly repressed in knockout (KO) mice when compared to that in WT mice of both models (Figure 1C; Supplementary Figure 1A). And this phenotype was further corroborated by decreased tumor weight at the endpoint of each experiment (Figure 1D; Supplementary Figure 1B).

**Figure 1.**
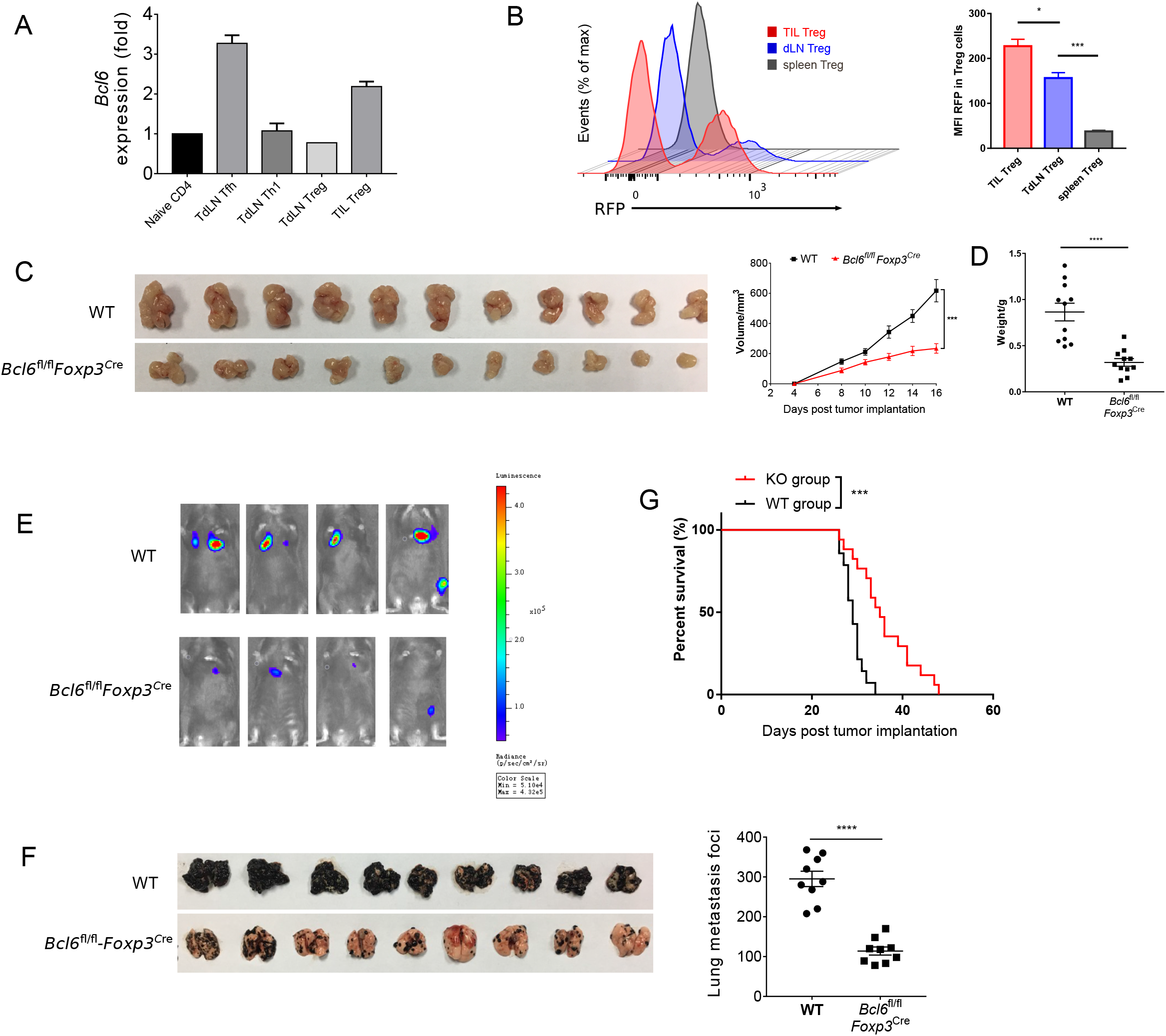
The deficiency of Bcl6 in Treg cells leads to delayed tumor progression. (**A**) RT-qPCR of Bcl6 expression in tumor infiltrating Treg (CD4^+^CD25^+^GITR^+^CXCR5^−^, noted as TIL Treg) cells, Tfh (gated in CD4^+^CD44^+^Foxp3^−^CXCR5^+^) and Th1 (gated in CD4^+^CD44^+^Foxp3^−^ CXCR5^−^) cells sorted from draining lymph node of WT tumor bearing mice. Naive CD4 (gated in CD4^+^CD44^−^) sorted from naive WT mice are included as control. Mice were inoculated with 5×10^5^ B16-F10 cells per mice intravenously, and were sacrificed (indicated as “Sac” in figure) at day 16 for analysis. The data presented are representative of two technical replicates pooled from at least 3 mice per group. (**B**) Bcl6 expression in tumor infiltrating Treg, dLNs Treg and spleen Treg (CD4^+^CD25^+^) cells. Bcl6-RFP reporter mice were inoculated with 1×10^6^ MC-38 subcutaneously, and were sacrificed at day 10 for flow cytometry analysis. Statistical analysis shown on the right. The data presented are representative of two independent experiments with at least 3 mice per group. Error bars indicate the mean±SEM. *p < 0.05, ***p < 0.001. (**C-D**) Pictures of MC-38 tumor samples harvested at day 17 after tumor implantation (1×10^6^ MC-38 cells per mice subcutaneously, C), with tumor growth curve and weight (D). (**E**) Tumor progression screened by bioluminescent imaging at D11. Mice were inoculated with 5×10^5^ B16-F10 with luciferase sequence intravenously per mice. (**F**) Pictures of lung samples harvested from metastasis model, in which mice were inoculated with 1×10^6^ B16-F10 cells per mice intravenously and sacrificed at day 16, with foci numbers calculated. (**G**) Survival curve of *Bcl6*^fl/fl^*Foxp3*^Cre^ and WT mice inoculated with 5×10^5^ B16-F10 per mice intravenously. Data were collected from 8-14 mice per group with two independent experiments. Statistical differences were calculated by unpaired t test (tumor weight), Log-rank test (survival curve) or two-way ANOVA with a post hoc Turkey test (tumor growth curve). ***p < 0.001, ****p < 0.0001. Data are presented as mean±SEM. See also Supplementary Figure 1

Next, we sought to investigate the role of Bcl6 during tumor metastasis. We challenged *Bcl6*^fl/fl^*Foxp3*^Cre^ and WT mice with B16-F10 melanoma cells intravenously to generate lung metastasis model. In order to continuously monitor tumor metastasis, we introduced the B-Luciferase B16-F10 cell line, which carries a luciferase-coding sequence in genome. At day 11 (D11) post tumor inoculation, we noted weaker fluorescent dots in KO mice with in-vivo imaging system (Figure 1E), indicating less metastatic foci in KO group than in control group. Consistently, we also observed less metastatic foci in KO mice compared to WT mice at day 18 post tumor inoculation (Figure 1F). In line with this result, *Bcl6*^fl/fl^*Foxp3*^Cre^ mice also demonstrated better survival rate when compared to WT counterparts (Figure 1G). Moreover, when proliferation-indicating dye labeled B16-F10 cells were intravenously transferred into WT and KO mice respectively, we harvested the same number of violet positive B16-F10 cells from lung tissues of both groups two hours later, indicating the comparable migration capacity of tumor cells within both groups. However, 20 hours later, a significant decrease in the number of pulmonary B16-F10 cells suggested that the lungs of KO mice represent a hostile environment for tumor cell engraftment (Supplementary Figure 1C).

### 3.2 The deficiency of Bcl6 impaired the suppressive capacity of Treg cells in TME

To better understand the role of Bcl6 in Treg cells during tumorigenesis, we examined tumor infiltrated effector T cell response after challenging *Bcl6*^fl/fl^*Foxp3*^Cre^ and WT mice with MC-38 cells subcutaneously. At D16 post tumor implantation, using flow cytometry analysis, we found that the deficiency of Bcl6 results in an impairment of Treg cells, evidenced by the decreased proportion and absolute number of Foxp3^+^CD4^+^ T cells (Figure 2A). Correspondingly, we observed increased tumor infiltrated CD4^+^CD44^+^ and CD8^+^CD44^+^ T cells upon Bcl6 deletion in Treg cells (Figure 2B). In intracellular cytokine staining, we found more tumor infiltrating CD8^+^CD44^+^ T cells generating CD107 and Interferon-gama (IFNγ) in KO mice compared to WT mice when stimulated with PMA plus Ionomycin (Figure 2C). Consistently, in melanoma lung metastasis model, we found a higher ratio of effector CD4 T cells to Treg cells in tumor compared with WT counterparts (Supplementary Figure 1D). Although the total number of CD8 T cells remain comparable between these two groups (Supplementary Figure 1D), CD8 T cells derived from KO mice did show better effector function than that of WT mice, shown by increased proportion and absolute number of CD107^+^, TNFα^+^ and KLRG1^+^ CD8^+^CD44^+^ T cells in KO mice (Supplementary Figure 1E). We also observed increased number of total CD4^+^CD44^+^ T cells and cytokine producing CD4^+^CD44^+^Foxp3^−^ T (CD107^+^ and TNFα^+^) cells (Supplementary Figure 1F). Thus, these data indicate the critical role of Bcl6 in regulating Treg cells’ suppression on effector T cells during tumorigenesis.

**Figure 2.**
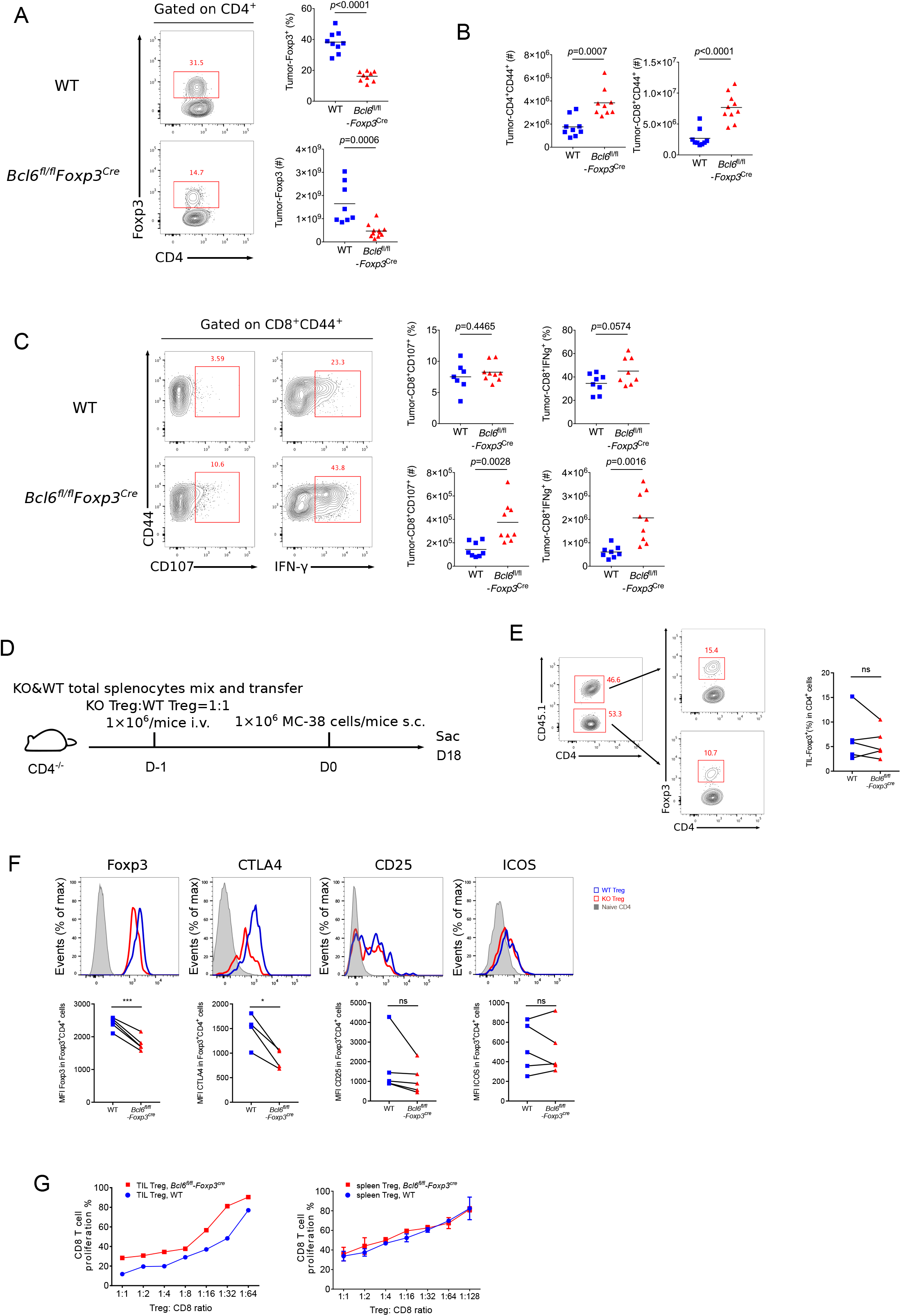
The deficiency of Bcl6 impaired the suppressive capacity of Treg cells. (**A-C**) Flow cytometry analysis, assessed at day 16 after MC-38 inoculation subcutaneously. The proportion and total number of Treg cells (A) and cytokine producing CD8 subsets (C). Summary of the total number of CD4^+^CD44^+^ and CD8^+^CD44^+^ T cells (B). The data (A-C) presented are representative of two independent experiments with 8-14 mice per group. # represents the absolute number and % represents the proportion of indicating population. Data are presented as mean±SEM. Unpaired t test. (**D**) Set up of splenic chimera mice, in situ model. Lymphocytes from WT and *Bcl6*^fl/fl^*Foxp3*^Cre^ mice were collected, mixed and transferred 1×10^6^ cells per mice to CD4^−/−^ recipients (Treg cells in a ratio of 1:1). On the following day, 1×10^6^ MC-38 cells were injected subcutaneously into recipient mice and mice were sacrificed at day 18 post tumor inoculation. (**E, F**) Flow cytometry of splenic chimera mice assay, analyzing the ratio of total CD4^+^T cells in KO versus WT mice, Treg cell proportion in CD4^+^ T cells (E), expression level of Foxp3, CTLA4, CD25 and ICOS in Treg cells (F). The data presented are representative of two independent experiments with at least 4 mice per group. Paired t test. *p < 0.05, ***p < 0.001, ns, no significant. (**G**) In vitro suppression assay. CD8^+^ T cell proliferation were assessed after incubated with tumor infiltrating Treg cells (CD4^+^CD25^+^GITR^+^, left) or naive spleen Treg cells (CD4^+^CD25^+^GITR^+^, right) sorted from *Bcl6*^fl/fl^*Foxp3*^Cre^ or WT mice in the ratio of 1:1 to 1:64 for 72 hours. See also Supplementary Figure 1 and S2.

To test whether Bcl6 play an intrinsic role in Treg cells, we generate a splenic chimera mice by mixing splenocytes derived from WT mice (CD45.1^+^, 50%) and KO mice (*Bcl6*^fl/fl^*Foxp3*^Cre^, CD45.2^+^, 50%) and injecting the mixtures into CD4^−/−^ recipients (Figure 2D). We compared the immune response of Treg cells derived from WT and KO donors in a same host of MC-38 primary tumor model. At the endpoint of tumor implantation, we observed comparable proportion of Treg cells originated from both donors (Figure 2E). However, KO mice derived Treg cells exhibited remarkably reduced expression of Foxp3 and CTLA4 in both tumor tissue (Figure 2F) and dLNs (Supplementary Figure 2A), suggesting a compromised stability and function of Bcl6-dificient Treg cells. And these phenomena were further corroborated in lung metastasis model (Supplementary Figure 2B). Consistently, we conducted in vitro suppression assay by co-culturing WT or Bcl6-deficient tumor infiltrating Treg cells with CFSE-labelled naive CD8 T cells that were followed by anti-CD3 and CD28 stimulation, and the results showed that Bcl6-deficient Treg cells exhibited impaired suppressive function compared with that of WT Treg cells. Contrarily, when isolating cells from naive mice, WT or Bcl6-deficient Treg cells exhibited comparable suppressive function (Figure 2G). Collectively, these data indicated that the abrogation of Bcl6 in Treg cells severely dampened the suppressive function of Treg cells in a cell intrinsic manner in TME, without profoundly influencing the absolute number of Treg cells.

### 3.3 Tfr cells are dispensable for tumor control

It has been reported that Bcl6 participates in regulating the differentiation of follicular regulatory T cells (Tfr) (33). which is a subset of Treg cells endowed with the capacity to inhibit germinal center reaction. To test whether tumor repression in KO mice was contributed by compromised Tfr response, we first examined Tfr population within tumor using CXCR5-GFP reporter mice (34). Notably, GFP^+^Foxp3^+^CD4^+^ T cells were barely detected in tumor, while a substantial population of Tfr was observed in dLNs (Supplementary Figure 2C), suggesting the absence of Tfr population in tumor mass. Consistent with published data (33), the deficiency of Bcl6 resulted in significantly decreased proportion and absolute number of Tfr cells in dLNs, accompanied with upregulated effector CD4 T cells including Th1 and Tfh cells (Supplementary Figure 2D). To investigate whether dLNs derived Tfr cells were involved in tumor control, we set up BM chimera mice by mixing bone marrow cells derived from *Foxp3*^DTR^ mice (50%) and *Cxcr5*^−/−^ mice (50%) and injecting the mixtures into sub-lethally irradiated CD4^−/−^ recipients (Figure 3A). In this setting, Tfr cells were selectively depleted after diphtheria (DT) administration (37), while Treg cells remained intact (Figure 3C). At D16 after tumor implantation (lung metastasis model), we found comparable tumor control ability between Tfr-deficient mice (DT treatment group) and control mice, indicated by similar lung metastasis foci between two groups (Figure 3B). Besides, DT induced deletion of Tfr cells did not influence tumor infiltrating Treg cells as well as effector CD4 T and CD8 T cells, evidenced by comparable number of Treg cells, Foxp3^−^CD44^+^CD4^+^ T (Figure 3D) and CD44^+^CD8^+^ T cells (Figure 3E). Thus, these data demonstrated that Tfr cells were dispensable for tumor control.

**Figure 3.**
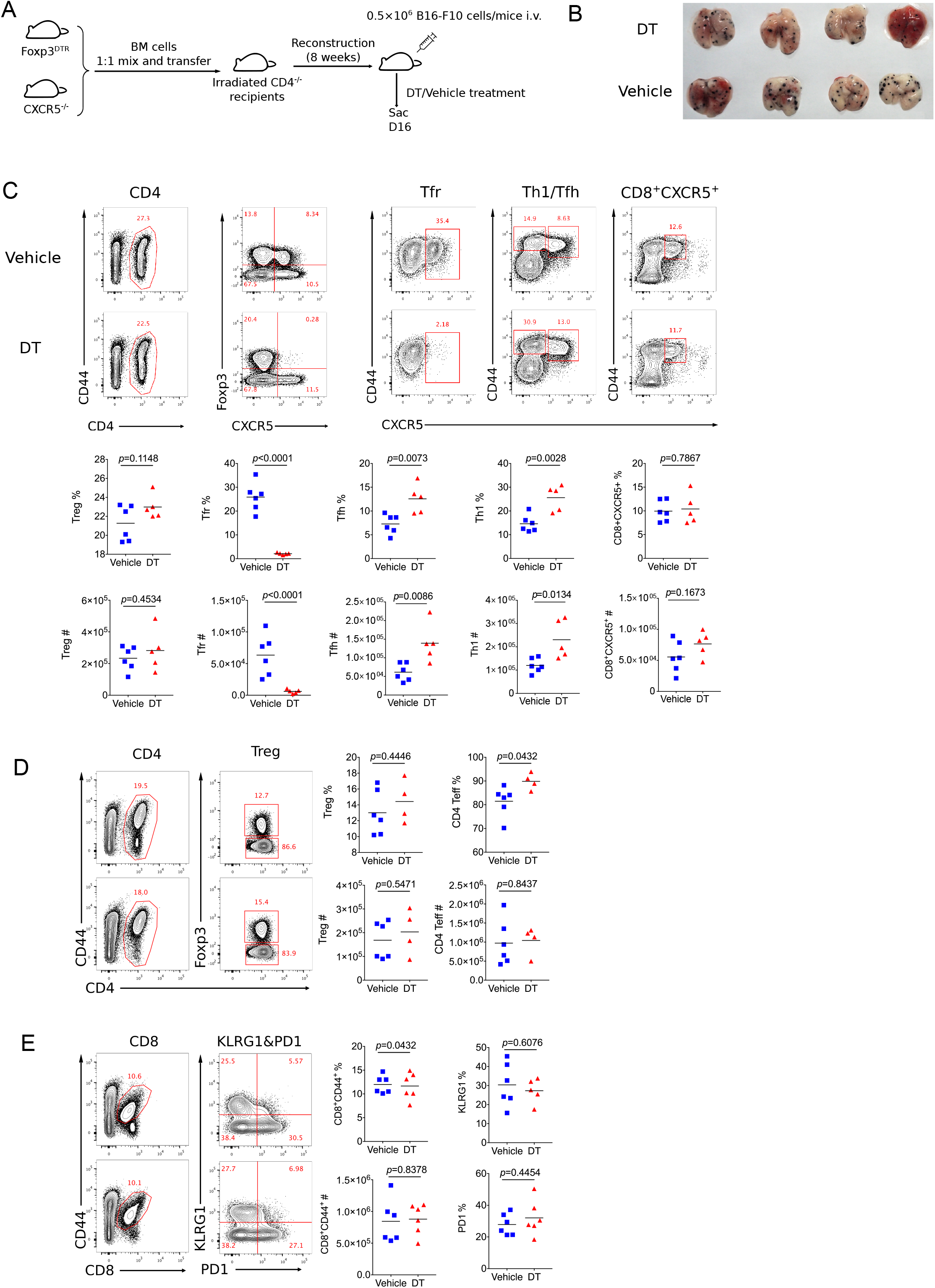
Tfr cells are dispensable for tumor control. (**A**) Set up of Tfr deficient Bone Marrow Chimera mice. Bone marrow cells were isolated from *Foxp3*^DTR^ and *Cxcr5*^−/−^ mice were mixed 1:1 and then transferred 8×10^6^ per mice to irradiated CD4^−/−^ mice. After reconstruction, mice were inoculated with 5×10^5^ B16-F10 cells intravenously and were treated with DT or vehicle, then sacrificed at D16 post tumor inoculation. (**B**) Pictures of lung samples harvested from metastasis model of Tfr deficient mice. (**C-E**) Flow cytometry analyzing proportion and total number of Treg, Tfr, Tfh, Th1, CD8^+^CXCR5^+^ subpopulation from dLNs (C) and Treg, CD4^+^CD44^−^ (D), CD8^+^ T cells (E) from tumor tissue. The data presented are representative of two independent experiments with at least 4 mice per group. # represents the absolute number and % represents the proportion of indicating population. Statistical differences were calculated by unpaired t test. Center values indicate mean. See also Supplementary Figure 2.

### 3.4 Bcl6 is essential in maintaining the lineage stability of Treg cells in TME

In order to illustrate the underlying mechanisms of Bcl6 in regulating Treg cell response during tumorigenesis, we sorted antigen experienced Treg, Th1 and Tfh cells as well as naive Treg cells from WT and KO mice at 16 days after tumor implantation to extract total RNA and conduct an array of quantitative real-time PCR (RT-qPCR) to profile the transcriptional signatures of the indicated subsets of CD4 T cells. Consistent with FACS data (shown in Figure 2F), the deficiency of Bcl6 resulted in reduced expression of Foxp3 in Treg cells in tumor model (Figure 4A). However, it had no effects on Foxp3 expression in naive state (Figure 4B), indicating that the identity of Bcl6-deficient Treg cells might be preferentially altered within TME. Accordingly, compared with WT counterparts, KO derived tumor infiltrating Treg cells upregulated the expression level of *Gata3*, which encodes the master transcriptional factor (TF) of Th2, as well as Th2 related cytokine IL4 (Figure 4C). Meanwhile, the expression of Th17 related cytokine IL17 also increased in KO mice despite similar level of retinoic acid receptor-related orphan receptor (ROR)γt between WT and KO group (Figure 4D). During naive state, however, KO Treg cells exhibited normal stability indicated by comparable level of *Gata3*, *Il17* and *Rorc* expression compared with WT mice (Figure 4E). These data suggested that the altered identity of Treg cells in KO mice was specifically confined to tumor micro-environment. Moreover, by analyzing Treg cells derived from different donors in spleen chimera mice (lung metastasis model), we further confirmed the enhanced expression of Tbet and GATA3 in KO Treg cells compared with WT ones in both tumor tissue (Figure 4F) and dLNs (Supplementary Figure 2E). All together, these data suggested the important role of Bcl6 in maintaining the lineage stability of Treg cells during tumorigenesis.

**Figure 4.**
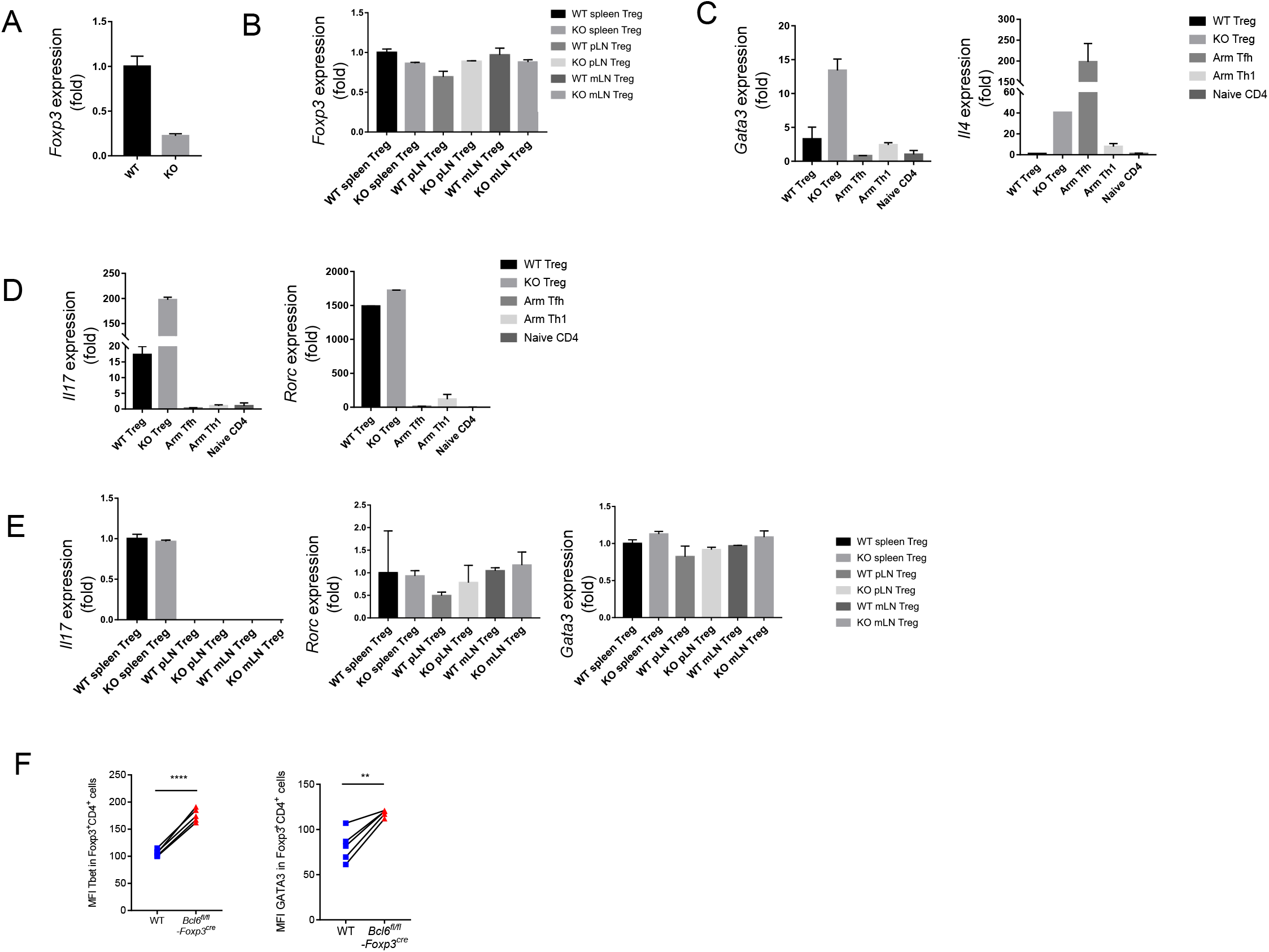
Bcl6 is essential in maintaining the lineage stability of Treg cells in TME. (**A**) RT-qPCR analysis of *Foxp3* expression, normalized to its expression in Bcl6-deficient tumor infiltrating Treg cells (CD4^+^CD25^+^GITR^+^CXCR5^−^), assessed at day 16 after B16-F10 inoculation intravenously. (**B**) RT-qPCR analysis of *Foxp3* expression, normalized to its expression in Treg cells (CD4^+^CD25^+^GITR^+^CXCR5^−^) from spleen, mesenteric lymph nodes(mLN) or peripheral non-draining lymph nodes (pLN) of naive *Bcl6*^fl/fl^*Foxp3*^Cre^ and WT mice. (**C, D**) RT-qPCR analysis of *Gata3* and *Il4* (c), *Il17* and *Rorc* (d) normalized to their expression in Bcl6-deficient and WT tumor infiltrating Treg cells (CD4^+^CD25^+^GITR^+^CXCR5^−^). Naive CD4 (gated in CD4^+^CD44^−^) sorted from naive WT mice, Tfh (gated in CD4^+^CD44^+^Foxp3^−^CXCR5^+^) and Th1 (gated in CD4^+^CD44^+^Foxp3^−^CXCR5^−^) cells sorted from WT mice infected with Armstrong (Arm) at day 8 are included as control. (**E**) RT-qPCR analysis of *Il17*, *Rorc* and *Gata3* normalized to their expression in Treg cells (CD4^+^CD25^+^GITR^+^CXCR5^−^) from spleen, mesenteric lymph nodes(mLN) or peripheral non-draining lymph nodes (pLN) of naive *Bcl6*^fl/fl^*Foxp3*^Cre^ and WT mice. The data (A-E) presented are representative of two technical replicates pooled from at least 3 mice per group. Error bars indicate the mean±SEM. (**F**) Flow cytometry analysis of tumor infiltrating Treg cells in splenic chimera mice assay, assessed at day 18 after B16-F10 inoculation intravenously. Quantification of Tbet and GATA3 in Bcl6-deficient and WT Treg cells. The data presented are representative of two independent experiments with at least 4 mice per group. Statistical differences were calculated by paired t test. **p < 0.01, ****p < 0.0001.

### 3.5 Bcl6 expression in Treg cells may predict the prognosis of human colorectal cancer

Given that the deficiency of Bcl6 in Treg cells resulted in repressed tumor growth in mouse models, we further checked whether Bcl6 expression could be exploited as an indicator for the clinical prognosis of human cancer patients. Interestingly, analysis of the independent human colorectal cancer cohort from TCGA (The Cancer Genome Atlas) database demonstrated that patients with higher level of Bcl6 (in Foxp3^+^CD4^+^ cells) showed significantly poorer overall survival compared to those with a lower level (Figure 5A). Moreover, the expression level of Bcl6 in Treg cells negatively correlates with the pathological grading of colorectal cancer, indicated by progressively increased proportion of Bcl6^hi^ Treg cells from pathological stage I to stage IV (TNM stage) colorectal cancer patients (Figure 5B). In a skin melanoma cohort from TCGA database, we observed increased proportion of lymph node metastasis in patients with higher Bcl6 expression in Treg cells (Supplementary Figure 2F), indicating that high level of Bcl6 in Treg cells may correlate with increased risk of metastasis of melanoma.

**Figure 5.**
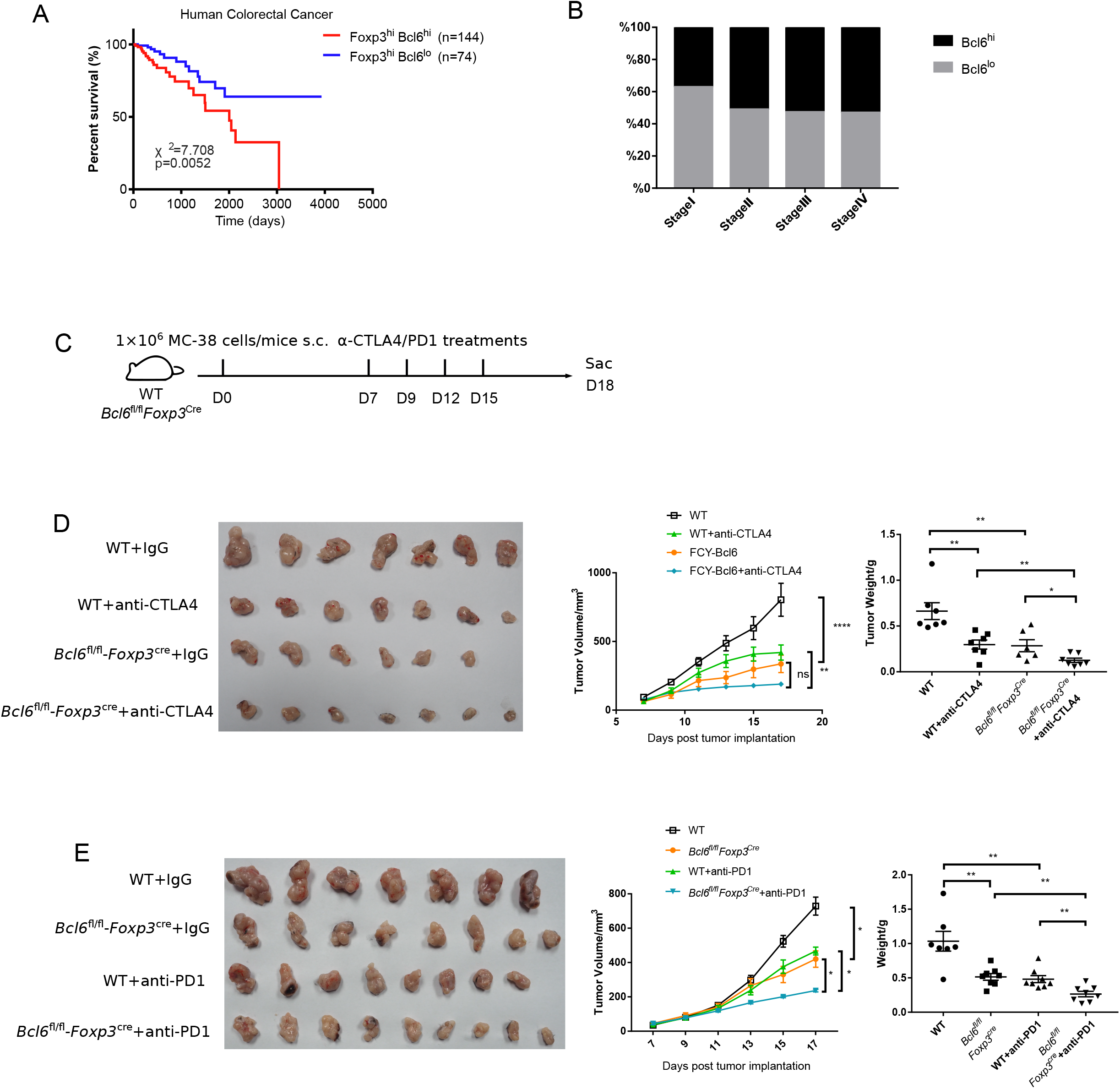
Bcl6 expression in Treg cells may predict the prognosis of human colorectal cancer. (**A**) Survival curves of Bcl6^hi^Foxp3^hi^ and Bcl6^lo^Foxp3^hi^ patients in human colorectal cancer TCGA dataset. (**B**) Proportion of Bcl6^hi^Foxp3^hi^ and Bcl6^lo^Foxp3^hi^ patients among Foxp3^hi^ patients in different stages of human colorectal cancer. (**C**) Set up of anti-CTLA4/anti-PD1 checkpoint blockade therapy. *Bcl6*^fl/fl^*Foxp3*^Cre^ and WT mice were implanted with MC-38 cells subcutaneously on day 0 and treated with anti-CTLA4 or anti-PD1 antibody at day 7, 9, 12 and 15. (**D, E**) Pictures of tumor samples harvested at day 18 after tumor implantation, with tumor weight and growth curve. Mice received either anti-CTLA4 (D) or anti-PD1 (E) therapy. The data were collected from 8-14 mice per group of two independent experiments. Statistical differences were calculated by unpaired t test (tumor weight) or two-way ANOVA with a post hoc Turkey test (tumor growth curve). *p < 0.05, **p < 0.01, ****p < 0.0001, ns, no significance. Data are presented as mean±SEM. See also Supplementary Figure 2.

Finally, we evaluated the synergetic effects of Bcl6 deletion with immune check point blockade (ICB). Consistent with published data (38–40), anti-CTLA4 or anti-PD1 alone significantly repressed tumor growth, which is comparable to that in KO group. Although Bcl6 deletion alone greatly improved the therapeutic efficacy, the combination of Bcl6 deletion and anti-CTLA4 or anti-PD1 did exhibit additional effects at delaying tumor growth (Figure 5C-E), indicating that Bcl6 can be a potential therapeutic target combined with ICB in cancer immunotherapy.

## 4 Discussion

Treg cells readily infiltrate into the TME and dampen anti-tumor immune responses, thereby becoming a barrier to effective cancer immunotherapy. In the past few years, Bcl6 has emerged as a central regulator of Tfh cell differentiation (35,41,42). However, the role of Bcl6 in Treg cells function has not been completely elucidated, particularly in terms of anti-tumor immune response. In this study, to specifically assess the role of Bcl6 in tumor infiltrating Treg cells, we challenged *Bcl6*^fl/fl^*Foxp3*^Cre^ mice with tumor cells and found that tumor growth was retarded in *Bcl6*^fl/fl^*Foxp3*^Cre^ mice with decreased tumor weight at the end point when compared to WT counterparts. And KO mice survive longer than WT mice in melanoma lung metastasis model. Besides, the deletion of Bcl6 in Treg cells resulted in impaired suppressive function, evidenced by enhanced functionality of CD8 T cells and non-Foxp3 CD4 T cells, as well as lowed expression of critical function markers of Treg cells including CTLA4 and CD25. Moreover, Bcl6 is essential in maintaining Treg cell lineage identity within tumor microenvironment. And tumor infiltrating *Bcl6*^−/−^ Treg cells exhibited markedly decreased expression of Foxp3, while upregulating the expression of TFs and cytokines related to other lineages, including Tbet, RORγt, GATA3, IL4 and IL17, suggesting a weakened lineage stability. Besides, Bcl6 expression in Treg cells reversely correlates with the prognosis of human colorectal cancer, while positively correlated with lymph node metastasis of skin melanoma. moreover, the deletion of Bcl6 in Treg cells acts coordinately with anti-CTLA4 or anti-PD1 to promote ICB therapeutic efficacy.

It is worth of noting that the impairment of the suppressive function of Treg cells induced by Bcl6 deficiency was only observed in tumor, while undetectable in naive state or non-draining lymph nodes (data not shown), which may be explained by very low expression levels of Bcl6 in Treg cells in this condition. Bcl6 conditional KO mice did not show uncontrolled inflammatory or autoimmune diseases at steady state. Additionally, compared to WT Treg cells, Bcl6-deficient ones in naive state exhibited normal expression of Foxp3 and comparable capacity in controlling T cell proliferation. Hence, it is important to investigate the specific role of Bcl6 in Treg cells during tumorigenesis. We hypothesize that the functional state and stability of tumor infiltrating Treg cells has been altered to a state in which Bcl6 is essentially required. Recent studies further distinguish peripheral Treg cells into resting Treg cells (TCF1^+^LEF1^+^CD62L^+^CD44^lo^), activated Treg cells (TCF1^+^LEF1^+/lo^CD62L^−^CD44^int/hi^) and effector Treg cells (TCF1^−^LEF1^−^CD62L^−^CD44^hi^) (43). Notably, Bcl6 was selectively enriched in activated Treg cells (43). In naive state, Treg cells contain a large fraction of resting Treg subsets, while only a small proportion of Treg cells are active or effector subsets. In tumor microenvironment, however, the highly suppressive signatures of Treg cells suggest an enrichment of activated or effector Treg subsets. Thus, Bcl6 may have a more pronounced effect in Treg cells within tumors.

Treg cells function is highly orchestrated, such that specific transcription factors regulate the ability of Treg cells to inhibit discrete types of T cell responses. It has been reported that T-bet uniquely controls the ability of Treg cells to suppress Th1 responses (44), IRF4 regulates the ability of Treg cells to suppress Th2 responses (45), and Stat3 directs the ability of Treg cells to suppress Th17 responses (46). Accordingly, Bcl6, which is the master regulator of Tfh cells, regulates the differentiation of Treg cells to Tfr and thus influence the ability of treg cells to control Tfh response (33). In this study, we found Bcl6 can also regulate Treg cell’s function in regulating the antitumor effects of other T cell lineages, which was shown in a Tfr independent pathway.

Bcl6 deletion significantly down regulate the expression of Foxp3 in tumor infiltrating Treg cells, which critically specifies the Treg lineage and maintains its functional program (47). But how Bcl6 regulate Foxp3 expression and function of tumor infiltrating Treg cells remains unsolved in our study. In Th cells, Bcl6 was reported to inhibit Th1, Th2, and Th17 development by suppressing Tbet, GATA3, and RORγt expression and activities (29,41,48). Meanwhile, it was reported that Bcl6 represses GATA3 expression in Treg cells and thus prevents Treg cells from acquiring Th2 effector-like characteristics. Consistently, we found an upregulation of Tbet, GATA3, and Th17 related cytokine (IL17) in KO Treg cells during tumorigenesis. However, whether Bcl6 uniquely targets these genes in Treg cells in this scenario requires further investigation.

Given the initial clinical successes achieved using PD1/PDL1 or CTLA4 antibody treatments in patients with cancer, the number of immunotherapy agents in clinical development is expanding rapidly with goals of improving the limited response rate and generating more durable responses (49,50). However, the clinical efficacy of ICBs is limited to only a small portion of patients, suggesting the importance to design new combinatory strategies (51). Currently, Treg cell targeted anti-cancer therapy have been tested clinically and/or preclinically. Some of the direct strategies involve targeting of those molecules specifically expressed by Treg cells, such as CTLA4, GITR, CD25 or ICOS. Several Treg cell-targeted therapies are under investigation alone or in combination with ICBs (1,52,53). Indeed, the anti-CTLA4 mAb ipilimumab can reinvigorate exhausted CD8 T cells while reportedly depleting Treg cells from the TME, which might contribute to the clinical benefits of this agent (54–57). Yet, one possible concern with Treg cell-targeted therapy is that systemic Treg cell depletion might considerably increase the risk of immune-related adverse events, such as autoimmunity owing to systemic disruption of immune tolerance (1,52). And several groups did observe an increased risk of colitis, rash and liver toxicity during the combined therapy of nivolumab with ipilimumab (an agent that can reportedly deplete or suppress Treg cells) (58,59). Mogamulizumab has also been associated with the development of rash and liver toxicity (60). For this reason, targeting Treg cells locally within the tumor could be a better approach that might avoid adverse events. In our study, we found that the induction of Bcl6 expression was strictly confined to tumor-infiltrating Treg cells, and the deletion of Bcl6 in Treg cells specifically repressed the function of this population, leaving systemic Treg cells (outside of TME) that do not express Bcl6 intact. Moreover, we demonstrated that targeting Bcl6 in Treg cells is sufficient to inhibit tumor growth and further improve the efficacy of ICB. This result reminds us that Bcl6 might be an optimal therapeutic candidate against cancer while reducing the side effects. Hence, an extensive and comprehensive understanding of Treg cell development, maintenance and function, especially in the scenario of tumorigenesis, could potentially lead to increases in the efficacy of Treg cell-targeted therapies and reduce the risk of adverse effects of such treatments.

Altogether, we found that Bcl6 is a negative regulator of the suppressive function of Treg cells during anti-tumor responses. The deletion of Bcl6 in Treg cells greatly improved the accumulation and function of effector T cells, unleashing a potent and synergistic therapeutic efficacy for immunotherapy.

## 5 Ethics Statement

All mouse experiments were performed in accordance with the guidelines of the Institutional Animal Care and Use Committees of the Third Military Medical University. This study was specifically reviewed and approved by the Laboratory Animal Welfare and Ethics Committee of the Third Military Medical University.

## 7 Author Contributions

YL, ZW, HL, LW, QL, QZ, JH, HW, JG, LX, JT, ZL, LH, QH, LX performed the experiments; XC analyzed TCGA data; LY designed the study, analyzed the data, and wrote the paper with QH, LX and YL; and LY supervised the study.

## 9 Acknowledgments

We thank Dr. Xindong Liu for generously providing us Bcl6-RFP mice. We thank Dr. Hua Tang for generously providing us *Foxp3*^DTR^ mice, and we thank the core facility center of Third Military Medical University for cell sorting.

## 1 Data Availability Statement

Publicly available datasets were analyzed in this study. These data can be found here: https://genome-cancer.ucsc.edu/

---

**Supplementary Figure 1. Related to Figure 1-2.**

(**A-B**) Pictures of B16-F10 tumor samples harvested on day 21 after tumor implantation (1×10^6^ B16-F10 cells per mice subcutaneously, A), with tumor growth curve and weight (B). (**C**) Set up and analysis of violet labeling assay. *Bcl6*^fl/fl^*Foxp3*^Cre^ and WT mice were inoculated with violet labeled B16-F10 cells intravenously per mice and were sacrificed 2 or 20 hours post tumor injection.

Summary of violet positive cells 2 or 20 hours post tumor injection are shown on the right. (**D-F**) Flow cytometry analysis assessed at day 16 after B16-F10 inoculation intravenously. Ratio of CD4^+^CD44^+^ and CD8^+^CD44^+^ T cells versus Treg cells, total number of CD4^+^CD44^+^ and CD8^+^CD44^+^ T cells (D). Proportion and total number of effecting (KLRG1^+^) as well as cytokine producing subsets (CD107^+^, TNFα^+^) of CD8^+^CD44^+^ T cells (E). Proportion and absolute number of cytokine-producing subsets of CD4^+^CD44^+^Foxp3^−^ cells (F). The data presented are representative of two independent experiments with 8-14 mice (D-F) or at least 4 mice (C) per group. # represents the absolute number and % represents the proportion of indicating population. Center values indicate mean. Unpaired t test.

**Supplementary Figure 2. Related to Figure 2, 3 and 5.**

(**A-B, E**) Flow cytometry of splenic chimera mice (B16-F10, metastasis model). Splenocytes from WT and *Bcl6*^fl/fl^*Foxp3*^Cre^ mice were collected and mixed with Treg cells in a ratio of 1:1, and were adoptively transferred (1×10^6^ cells per mice) to CD4^−/−^ recipients. On the following day, 1×10^6^ B16-F10 cells were injected intravenously into recipient mice and mice were sacrificed at day 18 post tumor inoculation. Quantification of Foxp3, CTLA4, CD25 and ICOS in Treg cells from dLNs (A) and tumor tissue (B). Quantification of Tbet and GATA3 in Treg cells from dLNs (E). The data presented are representative of two independent experiments with at least 4 mice per group. Paired t test. *p < 0.05, **p < 0.01, ***p < 0.001, ****p < 0.0001, ns, no significant. (**C**) Flow cytometry analysis of Tfr (gated in CD4^+^CD25^+^GITR^+^GFP^+^) cells in dLNs and tumor tissues isolated from CXCR5-GFP reporter mice, assessed at day 10 after MC-38 inoculation in situ. The data presented are representative of two independent experiments with at least 3 mice per group. (**D**) Flow cytometry analyzing total number of lymph node residing Treg cells (gated in CD4^+^CD44^+^Foxp3^+^CXCR5^−^), Tfr cells (gated in CD4^+^CD44^+^Foxp3^+^CXCR5^+^), Tfh cells (gated in CD4^+^CD44^+^Foxp3^−^CXCR5^+^) and Th1 cells (gated in CD4^+^CD44^+^Foxp3^−^CXCR5^−^), assessed at day 16 after B16-F10 inoculation to WT and *Bcl6*^fl/fl^*Foxp3*^Cre^ mice intravenously. The data presented are representative of two independent experiments with 8-14 per group. # represents the absolute number. Unpaired t test. Center values indicate mean. (**F**) Proportion of Bcl6^hi^Foxp3^hi^ and Bcl6^lo^Foxp3^hi^ patients among Foxp3^hi^ patients in primary and lymph node metastasis patients.

## References

1. Togashi Y, Shitara K, Nishikawa H. Regulatory T cells in cancer immunosuppression — implications for anticancer therapy. Nat Rev Clin Oncol (2019) 16:356–371. doi:10.1038/s41571-019-0175-7

2. Sawant D V., Vignali DAA. Once a Treg, always a Treg? Immunol Rev (2014) 259:173–191. doi:10.1111/imr.12173

3. Wan YY, Flavell RA. Regulatory T-cell functions are subverted and converted owing to attenuated Foxp3 expression. Nature (2007) 445:766–770. doi:10.1038/nature05479

4. Hori S. Control of Regulatory T Cell Development by the Transcription Factor Foxp3. Science (80-) (2003) 299:1057–1061. doi:10.1126/science.1079490

5. Sakaguchi S, Wing K, Onishi Y, Prieto-Martin P, Yamaguchi T. Regulatory T cells: How do they suppress immune responses? Int Immunol (2009) 21:1105–1111. doi:10.1093/intimm/dxp095

6. Shevach EM. Mechanisms of Foxp3+ T Regulatory Cell-Mediated Suppression. Immunity (2009) 30:636–645. doi:10.1016/j.immuni.2009.04.010

7. Salomon B, Lenschow DJ, Rhee L, Ashourian N, Singh B, Sharpe A, Bluestone JA. B7/CD28 costimulation is essential for the homeostasis of the CD4+CD25+ immunoregulatory T cells that control autoimmune diabetes. Immunity (2000) 12:431–440. doi:10.1016/S1074-7613(00)80195-8

8. Collison LW, Chaturvedi V, Henderson AL, Giacomin PR, Guy C, Bankoti J, Finkelstein D, Forbes K, Workman CJ, Brown SA, et al. IL-35-mediated induction of a potent regulatory T cell population. Nat Immunol (2010) 11:1093–1101. doi:10.1038/ni.1952

9. Tanaka A, Sakaguchi S. Regulatory T cells in cancer immunotherapy. Cell Res (2017) 27:109–118. doi:10.1038/cr.2016.151

10. Jameson SC, Masopust D. Understanding Subset Diversity in T Cell Memory. Immunity (2018) 48:214–226. doi:10.1016/j.immuni.2018.02.010

11. Smigiel KS, Richards E, Srivastava S, Thomas KR, Dudda JC, Klonowski KD, Campbell DJ. CCR7 provides localized access to IL-2 and defines homeostatically distinct regulatory T cell subsets. J Exp Med (2014) 211:121–136. doi:10.1084/jem.20131142

12. Mempel TR, Marangoni F. Guidance factors orchestrating regulatory T cell positioning in tissues during development, homeostasis, and response. Immunol Rev (2019) 289:129–141. doi:10.1111/imr.12761

13. Wyss L, Stadinski BD, King CG, Schallenberg S, Mccarthy NI, Lee JY, Kretschmer K, Terracciano LM, Anderson G, Surh CD, et al. Affinity for self antigen selects Treg cells with distinct functional properties. Nat Immunol (2016) 17:1093–1101. doi:10.1038/ni.3522

14. Togashi Y, Nishikawa H. “Regulatory T Cells: Molecular and Cellular Basis for Immunoregulation,” in Assessment & Evaluation in Higher Education, 3–27. doi:10.1007/82_2017_58

15. Sugiyama D, Nishikawa H, Maeda Y, Nishioka M, Tanemura A, Katayama I, Ezoe S, Kanakura Y, Sato E, Fukumori Y, et al. Anti-CCR4 mAb selectively depletes effector-type FoxP3+CD4+ regulatory T cells, evoking antitumor immune responses in humans. Proc Natl Acad Sci (2013) 110:17945–17950. doi:10.1073/pnas.1316796110

16. Ghiringhelli F, Menard C, Puig PE, Ladoire S, Roux S, Martin F, Solary E, Le Cesne A, Zitvogel L, Chauffert B. Metronomic cyclophosphamide regimen selectively depletes CD4+CD25+ regulatory T cells and restores T and NK effector functions in end stage cancer patients. Cancer Immunol Immunother (2007) 56:641–648. doi:10.1007/s00262-006-0225-8

17. Wing K, Onishi Y, Prieto-Martin P, Yamaguchi T, Miyara M, Fehervari Z, Nomura T, Sakaguchi S. CTLA-4 control over Foxp3+ regulatory T cell function. Science (80-) (2008) 322:271–275. doi:10.1126/science.1160062

18. Ge Y, Domschke C, Stoiber N, Schott S, Heil J, Rom J, Blumenstein M, Thum J, Sohn C, Schneeweiss A, et al. Metronomic cyclophosphamide treatment in metastasized breast cancer patients: immunological effects and clinical outcome. Cancer Immunol Immunother (2012) 61:353–362. doi:10.1007/s00262-011-1106-3

19. Tada Y, Togashi Y, Kotani D, Kuwata T, Sato E, Kawazoe A, Doi T, Wada H, Nishikawa H, Shitara K. Targeting VEGFR2 with Ramucirumab strongly impacts effector/ activated regulatory T cells and CD8+ T cells in the tumor microenvironment. J Immunother Cancer (2018) 6:1–14. doi:10.1186/s40425-018-0403-1

20. Saito T, Nishikawa H, Wada H, Nagano Y, Sugiyama D, Atarashi K, Maeda Y, Hamaguchi M, Ohkura N, Sato E, et al. Two FOXP3+CD4+ T cell subpopulations distinctly control the prognosis of colorectal cancers. Nat Med (2016) 22:679–684. doi:10.1038/nm.4086

21. Curiel TJ, Coukos G, Zou L, Alvarez X, Cheng P, Mottram P, Evdemon-Hogan M, Conejo-Garcia JR, Zhang L, Burow M, et al. Specific recruitment of regulatory T cells in ovarian carcinoma fosters immune privilege and predicts reduced survival. Nat Med (2004) 10:942–949. doi:10.1038/nm1093

22. De Simone M, Arrigoni A, Rossetti G, Gruarin P, Ranzani V, Politano C, Bonnal RJP, Provasi E, Sarnicola ML, Panzeri I, et al. Transcriptional Landscape of Human Tissue Lymphocytes Unveils Uniqueness of Tumor-Infiltrating T Regulatory Cells. Immunity (2016) 45:1135–1147. doi:10.1016/j.immuni.2016.10.021

23. Plitas G, Konopacki C, Wu K, Bos PD, Morrow M, Putintseva E V., Chudakov DM, Rudensky AY. Regulatory T Cells Exhibit Distinct Features in Human Breast Cancer. Immunity (2016) 45:1122–1134. doi:10.1016/j.immuni.2016.10.032

24. Czerwinska P, Rucinski M, Wlodarczyk N, Jaworska A, Grzadzielewska I, Gryska K, Galus L, Mackiewicz J, Mackiewicz A. Therapeutic melanoma vaccine with cancer stem cell phenotype represses exhaustion and maintains antigen-specific T cell stemness by up-regulating BCL6. Oncoimmunology (2020) 9:1710063. doi:10.1080/2162402X.2019.1710063

25. Huang Q, Xu L, Ye L. T cell immune response within B-cell follicles. 1st ed. Elsevier Inc. (2019). doi:10.1016/bs.ai.2019.08.008

26. Sage PT, Sharpe AH. T follicular regulatory cells. Immunol Rev (2016) 271:246–259. doi:10.1111/imr.12411

27. Vinuesa CG, Linterman MA, Yu D, MacLennan ICM. Follicular Helper T Cells. Annu Rev Immunol (2016) 34:335–368. doi:10.1146/annurev-immunol-041015-055605

28. Liu X, Yan X, Zhong B, Nurieva RI, Wang A, Wang X, Martin-Orozco N, Wang Y, Chang SH, Esplugues E, et al. Bcl6 expression specifies the T follicular helper cell program in vivo. J Exp Med (2012) 209:1841–1852. doi:10.1084/jem.20120219

29. Nurieva RI, Chung Y, Martinez GJ, Yang XO, Tanaka S, Matskevitch TD, Wang Y-H, Dong C. Bcl6 Mediates the Development of T Follicular Helper Cells. Science (80-) (2009) 325:1001–1005. doi:10.1126/science.1176676

30. Rawal S, Fan H, Liu Z, Lim H, Tanaka S, Neelapu SS, Chu F, Wang Y-H, Nurieva RI, Martinez GJ, et al. Follicular regulatory T cells expressing Foxp3 and Bcl-6 suppress germinal center reactions. Nat Med (2011) 17:983–988. doi:10.1038/nm.2426

31. Sawant D V., Wu H, Yao W, Sehra S, Kaplan MH, Dent AL. The transcriptional repressor Bcl6 controls the stability of regulatory T cells by intrinsic and extrinsic pathways. Immunology (2015) 145:11–23. doi:10.1111/imm.12393

32. Sawant D V., Sehra S, Nguyen ET, Jadhav R, Englert K, Shinnakasu R, Hangoc G, Broxmeyer HE, Nakayama T, Perumal NB, et al. Bcl6 Controls the Th2 Inflammatory Activity of Regulatory T Cells by Repressing Gata3 Function. J Immunol (2012) 189:4759–4769. doi:10.4049/jimmunol.1201794

33. Xu L, Huang Q, Wang H, Hao Y, Bai Q, Hu J, Li Y, Wang P, Chen X, He R, et al. The Kinase mTORC1 Promotes the Generation and Suppressive Function of Follicular Regulatory T Cells. Immunity (2017) 47:538–551.e5. doi:10.1016/j.immuni.2017.08.011

34. He R, Hou S, Liu C, Zhang A, Bai Q, Han M, Yang Y, Wei G, Shen T, Yang X, et al. Follicular CXCR5-expressing CD8+ T cells curtail chronic viral infection. Nature (2016) 537:412–428. doi:10.1038/nature19317

35. Xu L, Cao Y, Xie Z, Huang Q, Bai Q, Yang X, He R, Hao Y, Wang H, Zhao T, et al. The transcription factor TCF-1 initiates the differentiation of TFH cells during acute viral infection. Nat Immunol (2015) 16:991–999. doi:10.1038/ni.3229

36. Johnston RJ, Poholek AC, DiToro D, Yusuf I, Eto D, Barnett B, Dent AL, Craft J, Crotty S. Bcl6 and Blimp-1 Are Reciprocal and Antagonistic Regulators of T Follicular Helper Cell Differentiation. Science (80-) (2009) 325:1006–1010. doi:10.1126/science.1175870

37. Botta D, Fuller MJ, Marquez-Lago TT, Bachus H, Bradley JE, Weinmann AS, Zajac AJ, Randall TD, Lund FE, León B, et al. Dynamic regulation of T follicular regulatory cell responses by interleukin 2 during influenza infection. Nat Immunol (2017) 18:1249–1260. doi:10.1038/ni.3837

38. Ngiow SF, Young A, Jacquelot N, Yamazaki T, Enot D, Zitvogel L, Smyth MJ. A Threshold Level of Intratumor CD8 + T-cell PD1 Expression Dictates Therapeutic Response to Anti-PD1. Cancer Res (2015) 75:3800–3811. doi:10.1158/0008-5472.CAN-15-1082

39. Ngwa W, Irabor OC, Schoenfeld JD, Hesser J, Demaria S, Formenti SC. Using immunotherapy to boost the abscopal effect. Nat Rev Cancer (2018) 18:313–322. doi:10.1038/nrc.2018.6

40. Selby MJ, Engelhardt JJ, Quigley M, Henning KA, Chen T, Srinivasan M, Korman AJ. Anti-CTLA-4 antibodies of IgG2a isotype enhance antitumor activity through reduction of intratumoral regulatory T cells. Cancer Immunol Res (2013) 1:32–42. doi:10.1158/2326-6066.CIR-13-0013

41. Yu D, Rao S, Tsai LM, Lee SK, He Y, Sutcliffe EL, Srivastava M, Linterman M, Zheng L, Simpson N, et al. The Transcriptional Repressor Bcl-6 Directs T Follicular Helper Cell Lineage Commitment. Immunity (2009) 31:457–468. doi:10.1016/j.immuni.2009.07.002

42. Nurieva RI, Chung Y, Martinez GJ, Yang XO, Tanaka S, Matskevitch TD, Wang Y-H, Dong C. Bcl6 mediates the development of T follicular helper cells. Science (2009) 325:1001–5. doi:10.1126/science.1176676

43. Yang B-H, Wang K, Wan S, Liang Y, Yuan X, Dong Y, Cho S, Xu W, Jepsen K, Feng G-S, et al. TCF1 and LEF1 Control Treg Competitive Survival and Tfr Development to Prevent Autoimmune Diseases. Cell Rep (2019) 27:3629–3645.e6. doi:10.1016/j.celrep.2019.05.061

44. Koch MA, Tucker-Heard G, Perdue NR, Killebrew JR, Urdahl KB, Campbell DJ. The transcription factor T-bet controls regulatory T cell homeostasis and function during type 1 inflammation. Nat Immunol (2009) 10:595–602. doi:10.1038/ni.1731

45. Zheng Y, Chaudhry A, Kas A, DeRoos P, Kim JM, Chu T-T, Corcoran L, Treuting P, Klein U, Rudensky AY. Regulatory T-cell suppressor program co-opts transcription factor IRF4 to control TH2 responses. Nature (2009) 458:351–356. doi:10.1038/nature07674

46. Chaudhry A, Rudra D, Treuting P, Samstein RM, Liang Y, Kas A, Rudensky AY. CD4+ Regulatory T Cells Control TH17 Responses in a Stat3-Dependent Manner. Science (80-) (2009) 326:986–991. doi:10.1126/science.1172702

47. Fontenot JD, Rasmussen JP, Williams LM, Dooley JL, Farr AG, Rudensky AY. Regulatory T Cell Lineage Specification by the Forkhead Transcription Factor Foxp3. Immunity (2005) 22:329–341. doi:10.1016/j.immuni.2005.01.016

48. Mondal A, Sawant D, Dent AL. Transcriptional Repressor BCL6 Controls Th17 Responses by Controlling Gene Expression in Both T Cells and Macrophages. J Immunol (2010) 184:4123–4132. doi:10.4049/jimmunol.0901242

49. McLane LM, Abdel-Hakeem MS, Wherry EJ. CD8 T Cell Exhaustion During Chronic Viral Infection and Cancer. Annu Rev Immunol (2019) 37:457–495. doi:10.1146/annurev-immunol-041015-055318

50. Patel SA, Minn AJ. Combination Cancer Therapy with Immune Checkpoint Blockade: Mechanisms and Strategies. Immunity (2018) 48:417–433. doi:10.1016/j.immuni.2018.03.007

51. Hashimoto M, Kamphorst AO, Im SJ, Kissick HT, Pillai RN, Ramalingam SS, Araki K, Ahmed R. CD8 T Cell Exhaustion in Chronic Infection and Cancer: Opportunities for Interventions. doi:10.1146/annurev-med-012017-043208

52. Raffin C, Vo LT, Bluestone JA. Treg cell-based therapies: challenges and perspectives. Nat Rev Immunol (2019) doi:10.1038/s41577-019-0232-6

53. Zou W. Regulatory T cells, tumour immunity and immunotherapy. Nat Rev Immunol (2006) 6:295–307. doi:10.1038/nri1806

54. Vargas FA, Furness AJS, Litchfield K, Joshi K, Rosenthal R, Ghorani E, Solomon I, Lesko MH, Ruef N, Roddie C, et al. Fc Effector Function Contributes to the Activity of Human Anti-CTLA-4 Antibodies. Cancer Cell (2018) 33:649–663.e4. doi:10.1016/j.ccell.2018.02.010

55. Hodi FS, O’Day SJ, McDermott DF, Weber RW, Sosman JA, Haanen JB, Gonzalez R, Robert C, Schadendorf D, Hassel JC, et al. Improved Survival with Ipilimumab in Patients with Metastatic Melanoma. N Engl J Med (2010) 363:711–723. doi:10.1056/NEJMoa1003466

56. Bulliard Y, Jolicoeur R, Windman M, Rue SM, Ettenberg S, Knee DA, Wilson NS, Dranoff G, Brogdon JL. Activating fc γ receptors contribute to the antitumor activities of immunoregulatory receptor-targeting antibodies. J Exp Med (2013) 210:1685–1693. doi:10.1084/jem.20130573

57. Simpson TR, Li F, Montalvo-Ortiz W, Sepulveda MA, Bergerhoff K, Arce F, Roddie C, Henry JY, Yagita H, Wolchok JD, et al. Fc-dependent depletion of tumor-infiltrating regulatory t cells co-defines the efficacy of anti-CTLA-4 therapy against melanoma. J Exp Med (2013) 210:1695–1710. doi:10.1084/jem.20130579

58. Hellmann MD, Ciuleanu TE, Pluzanski A, Lee JS, Otterson GA, Audigier-Valette C, Minenza E, Linardou H, Burgers S, Salman P, et al. Nivolumab plus ipilimumab in lung cancer with a high tumor mutational burden. N Engl J Med (2018) 378:2093–2104. doi:10.1056/NEJMoa1801946

59. Larkin J, Chiarion-Sileni V, Gonzalez R, Grob JJ, Cowey CL, Lao CD, Schadendorf D, Dummer R, Smylie M, Rutkowski P, et al. Combined nivolumab and ipilimumab or monotherapy in untreated Melanoma. N Engl J Med (2015) 373:23–34. doi:10.1056/NEJMoa1504030

60. Kurose K, Ohue Y, Wada H, Iida S, Ishida T, Kojima T, Doi T, Suzuki S, Isobe M, Funakoshi T, et al. Phase Ia Study of FoxP3+ CD4 Treg Depletion by Infusion of a Humanized Anti-CCR4 Antibody, KW-0761, in Cancer Patients. Clin Cancer Res (2015) 21:4327–4336. doi:10.1158/1078-0432.CCR-15-0357

