## Supplementary Materials for "Bcl6 Preserves the Suppressive Function of Regulatory T Cells during Tumorigenesis"

### 1 Supplementary Figures and Tables

#### 1.1 Supplementary Figures

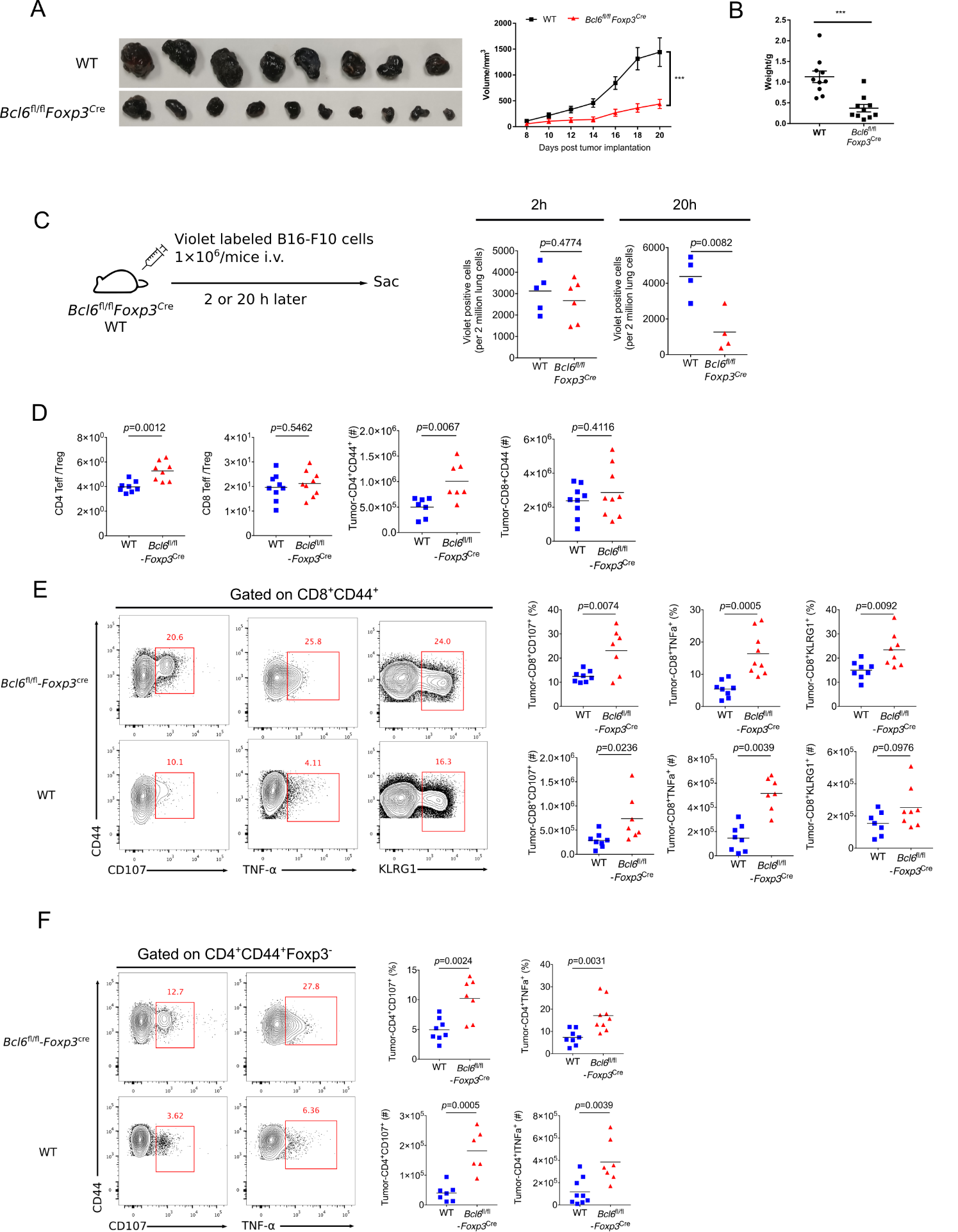

**Supplementary Figure 1.**

(**A-B**) Pictures of B16-F10 tumor samples harvested on day 21 after tumor implantation (1×10^6^ B16-F10 cells per mice subcutaneously, A), with tumor growth curve and weight (B). (**C**) Set up and analysis of violet labeling assay. *Bcl6*^fl/fl^*Foxp3*^Cre^ and WT mice were inoculated with violet labeled B16-F10 cells intravenously per mice and were sacrificed 2 or 20 hours post tumor injection. Summary of violet positive cells 2 or 20 hours post tumor injection are shown on the right. (**D-F**) Flow cytometry analysis assessed at day 16 after B16-F10 inoculation intravenously. Ratio of CD4^+^CD44^+^ and CD8^+^CD44^+^ T cells versus Treg cells, total number of CD4^+^CD44^+^ and CD8^+^CD44^+^ T cells (D). Proportion and total number of effecting (KLRG1^+^) as well as cytokine producing subsets (CD107^+^, TNFα^+^) of CD8^+^CD44^+^ T cells (E). Proportion and absolute number of cytokine-producing subsets of CD4^+^CD44^+^Foxp3^-^ cells (F). The data presented are representative of two independent experiments with 8-14 mice (D-F) or at least 4 mice (C) per group. # represents the absolute number and % represents the proportion of indicating population. Center values indicate mean. Unpaired t test.

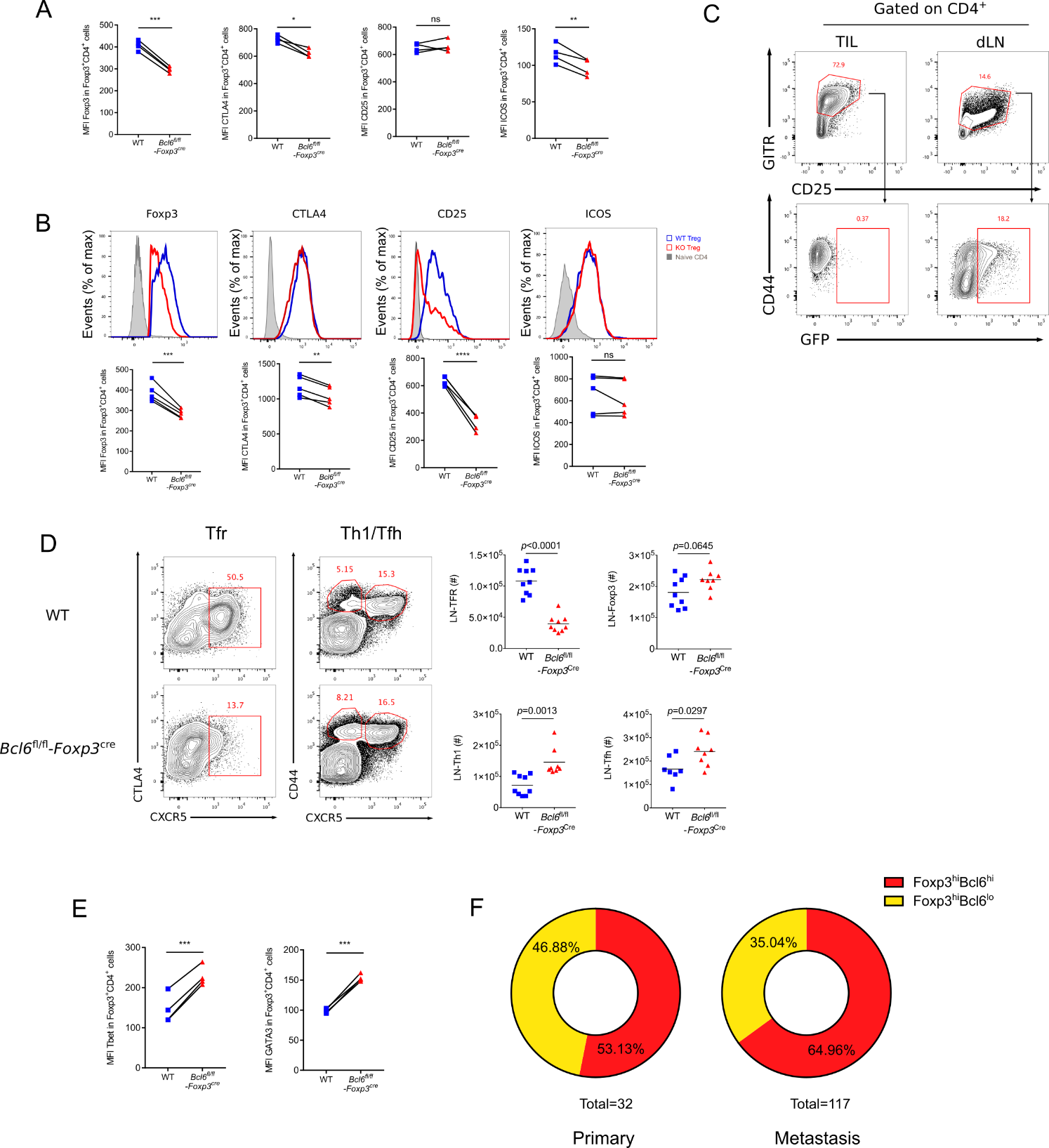

**Supplementary Figure 2.**

(**A-B, E**) Flow cytometry of splenic chimera mice (B16-F10, metastasis model). Splenocytes from WT and *Bcl6*^fl/fl^*Foxp3*^Cre^ mice were collected and mixed with Treg cells in a ratio of 1:1, and were adoptively transferred (1×10^6^ cells per mice) to CD4^-/-^ recipients. On the following day, 1×10^6^ B16-F10 cells were injected intravenously into recipient mice and mice were sacrificed at day 18 post tumor inoculation. Quantification of Foxp3, CTLA4, CD25 and ICOS in Treg cells from dLNs (A) and tumor tissue (B). Quantification of Tbet and GATA3 in Treg cells from dLNs (E). The data presented are representative of two independent experiments with at least 4 mice per group. Paired t test. *p < 0.05, **p < 0.01, ***p < 0.001, ****p < 0.0001, ns, no significant. (**C**) Flow cytometry analysis of Tfr (gated in CD4^+^CD25^+^GITR^+^GFP^+^) cells in dLNs and tumor tissues isolated from CXCR5-GFP reporter mice, assessed at day 10 after MC-38 inoculation in situ. The data presented are representative of two independent experiments with at least 3 mice per group. (**D**) Flow cytometry analyzing total number of lymph node residing Treg cells (gated in CD4^+^CD44^+^Foxp3^+^CXCR5^-^), Tfr cells (gated in CD4^+^CD44^+^Foxp3^+^CXCR5^+^), Tfh cells (gated in CD4^+^CD44^+^Foxp3^-^CXCR5^+^) and Th1 cells (gated in CD4^+^CD44^+^Foxp3^-^CXCR5^-^), assessed at day 16 after B16-F10 inoculation to WT and *Bcl6*^fl/fl^*Foxp3*^Cre^ mice intravenously. The data presented are representative of two independent experiments with 8-14 per group. # represents the absolute number. Unpaired t test. Center values indicate mean. (**F**) Proportion of Bcl6^hi^Foxp3^hi^ and Bcl6^lo^Foxp3^hi^ patients among Foxp3^hi^ patients in primary and lymph node metastasis patients.

#### 1.2 Supplementary Tables

**Supplementary Table 1. Antibodies and reagents used in flow cytometry**

| **Antibody target/Reagent** | **Clone** | **Provider** |
| --- | --- | --- |
| CD4 | RM4-5 | Biolegend |
| CD8 | 100734 | Biolegend |
| CD44 | IM7 | eBioscience |
| CD45.1 | A20 | Biolegend |
| CD45.2 | 104 | Biolegend |
| PD-1 | RMP1-30 | eBioscience |
| CXCR5 | 2G8 | BD Biosciences |
| Biotin Goat Anti-Rat IgG | 112-065-143 | Jackson Immunoresearch |
| Streptavidin | 25-4317-82 | eBioscience |
| Live/Dead Kit | L10119 | Life Technologies |
| Bcl2 | 7/Bcl-2 | BD Biosciences |
| Annexin V Kit | 88-8102 | eBioscience |
| BrdU | 3D4 | BD Biosciences |
| Ki-67 | B56 | BD Biosciences |
| Granzyme B | GB11 | Life technologies |
| IFNγ | XMG1.2 | BD Biosciences |
| TNFα | MP6-XT22 | BD Biosciences |
| CD107a | 1D48 | BD Biosciences |
| CD107b | ABL-93 | BD Biosciences |
| Bcl6 | K112-91 | BD Biosciences |
| Foxp3 | FJK-16s | eBioscience |
| CD25 | PC61.5 | Biolegend |
| ICOS | C398.4A | Biolegend |
| GITR | DTA-1 | eBioscience |
| CTLA4 | UC10-4B9 | Biolegend |

**Supplementary Table 2. Primers used in quantitative PCR**

| **Gene symbol** | **Forward primer** | **Reverse primer** |
| --- | --- | --- |
| ***Foxp3*** | CCCATCCCCAGGAGTCTTG | ACCATGACTAGGGGCACTGTA |
| ***Gata3*** | CTCGGCCATTCGTACATGGAA | GGATACCTCTGCACCGTAGC |
| ***Il4*** | TTACCTTGACGGTGTTCATACAG | TCTGCTCCTATTCGACCACTATC |
| ***Il17*** | CACCCCCGGAACACCAAAG | CATACTCTTCCATTCGAGCGTAG |
| ***Rorc*** | GACCCACACCTCACAAATTGA | AGTAGGCCACATTACACTGCT |
